# Expanding Targeted Instrumentation for Discovery Applications: Complement Reporter Ion Quantification with a Quadrupole-Ion Trap Instrument

**DOI:** 10.1101/2025.04.18.649483

**Authors:** Edward R. Cruz, Alexander N.T. Johnson, Vyas Pujari, Thao Nguyen, Michael Stadlmeier, Jessica Wang, Cristina Jacob, Graeme C. McAlister, Philip M. Remes, Martin Wühr

## Abstract

Proteomics workflows have traditionally been divided into discovery-based and targeted approaches, with instrumentation optimized specifically for each. Discovery experiments typically utilize high-resolution analyzers while targeted workflows rely on the sensitivity and specificity of triple quadrupole systems. Recently, a quadrupole-ion trap instrument (Stellar™ MS) demonstrated superior performance for targeted applications compared to conventional triple quadrupoles. In this study, we expand the capabilities of this platform to multiplexed shotgun proteomics using complement reporter ion quantification in an ion trap (iTMTproC). Benchmarking experiments with defined standards show that iTMTproC achieves quantification accuracy and interference reduction comparable to MultiNotch MS3 on the Orbitrap Fusion Lumos™, a dedicated quadrupole-ion trap-Orbitrap™ tribrid instrument optimized for this purpose. Notably, iTMTproC quantifies slightly more proteins than MultiNotch MS3. We further validate this approach through a developmental time-series analysis of frog embryos, obtaining proteomic data nearly indistinguishable from MultiNotch MS3, with slightly increased protein quantification depth. These findings significantly extend the functionality of targeted instrumentation, underscoring the versatility of quadrupole-ion trap systems and providing cost-effective access to highly accurate, multiplexed quantitative shotgun proteomics.

## Introduction

Proteomics experiments traditionally follow two distinct paradigms: discovery-based (shotgun) approaches and targeted analyses^1-4^. Discovery proteomics aims to identify and quantify thousands of proteins in complex samples, typically using high-resolution mass analyzers capable of resolving and identifying diverse peptide populations in an unbiased manner^5, 6^. In contrast, targeted proteomics measures predefined subsets of proteins or peptides, achieving higher quantitative precision and reproducibility across multiple samples^1, 2, 7^. Classic targeted workflows often utilize triple quadrupole or quadrupole–ion trap instruments for selected or multiple reaction monitoring (SRM/MRM), enabling rapid, sensitive measurements at nominal mass resolution^8-10^. Recently, a quadrupole–ion trap mass spectrometer (Stellar™ MS) demonstrated superior performance in targeted workflows compared to conventional triple-quadrupole instruments, achieving a 10-fold improvement in limits of quantitation and a 2-fold improvement in detection limits^11^ (Figure 1a-b).

**Figure 1.**
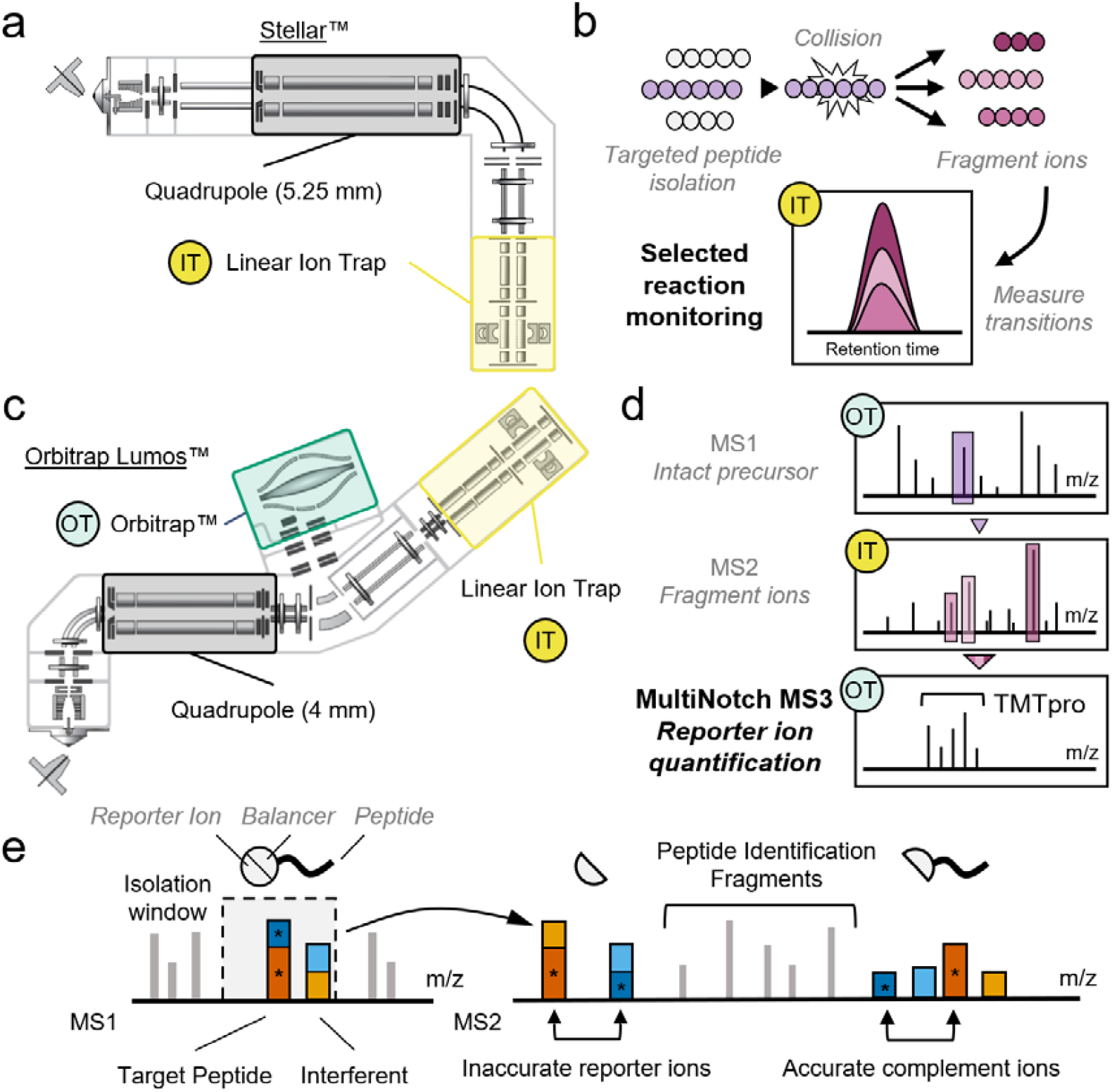
Comparison of Orbitrap Lumos™ and Stellar™ and their primary intended usage. **(a)** Schematic of the Stellar™. The Stellar™ is optimized for high-sensitivity targeted applications, incorporating a linear ion trap and a state-of-the-art quadrupole (field radius = 5.25 mm), which surpasses the quadrupole used in the Orbitrap Lumos™ (field radius = 4 mm). Analyzers shared with the Orbitrap Lumos™ are highlighted in black and yellow. In later subpanels, the Orbitrap™ is highlighted in green, and in the example workflows, OT (Orbitrap™) and IT (ion trap) indicate which analyzer performs the corresponding scan. **(b)** Targeted proteomics on the Stellar™. The Stellar™ is tailored for targeted workflows such as selected reaction monitoring (SRM), in which peptides of interest are selectively isolated, fragmented, and their fragment ions (“transitions”) are monitored for quantitative analysis. **(c)** Schematic of the Orbitrap Lumos™. The Orbitrap Lumos™ is a tribrid mass spectrometer designed to support accurate multiplexed shotgun proteomics using MultiNotch MS3. It integrates three mass analyzers: a quadrupole, an Orbitrap™, and a linear ion trap. **(d)** Workflow of MultiNotch MS3 multiplexed shotgun proteomics on the Orbitrap Lumos™. In MultiNotch MS3 using TMTpro™ reagents, synchronous precursor selection (SPS) is employed to isolate multiple fragment ions from the MS2 scan simultaneously. These selected ions are further fragmented in an MS3 scan to generate reporter ions for quantification. This additional fragmentation step reduces interference from co-isolated peptides, thereby improving quantification accuracy in complex samples. **(e)** Principle of complement reporter ion quantification (TMTproC). During MS1 isolation, the isolation window captures both the target peptide (dark orange and blue) and co-isolated interfering peptides (light orange and blue). In conventional MS2-based workflows, these co-isolated peptides produce indistinguishable reporter ions, leading to quantification errors. TMTproC addresses this by leveraging complement ions, which retain the intact peptide backbone along with the balancer group. Because this balancer–peptide conjugate encodes both peptide identity and sample origin, TMTproC reduces interference from unrelated peptides and enables more accurate quantification.

Among discovery-based proteomics techniques, two leading strategies are data independent acquisition (DIA) and multiplexed proteomics based on isobaric tagging^12-16^. Isobaric mass tags, such as TMTpro, enable the simultaneous quantification of peptides from multiple samples within a single experiment^17, 18^. This multiplexing approach substantially boosts throughput and experimental consistency by eliminating inter-run variability. However, early MS2-based implementations of multiplexing faced a key obstacle: reporter ion ratio distortion caused by co-isolated peptides, leading to inaccurate quantification^19-21^. To mitigate interference, MS3-based methods were introduced, utilizing an additional fragmentation step (MultiNotch MS3) to co-isolate and fragment b- and y-ions specifically related to the targeted peptide^19, 22^ (Figure 1c-d). While this MS3 strategy substantially reduces ratio distortion and enhances quantification accuracy, it sacrifices acquisition speed and requires specialized hybrid mass spectrometers, i.e. tribrid quadrupole–ion trap–Orbitrap™ instruments. Recent advancements incorporate real-time search (RTS) capabilities with MultiNotch MS3, allowing MS3 scans to be selectively triggered only upon successful peptide identifications during MS2 analysis^23, 24^. This innovation further enhances measurement accuracy by exclusively isolating b- and y-ions carrying isobaric tags from peptides of interest^24^.

An alternative strategy for accurate multiplexed quantification at the MS2 level, known as TMTproC, utilizes complementary reporter ions^25-28^. Unlike traditional low-mass reporter ion quantification, TMTproC leverages balancer-group fragment ions that retain isotopic label information in the mid- to high-mass region. Because complementary ion masses are precursor-specific, they substantially reduce co-isolation interference (Figure 1e). To date, TMTproC methods have been implemented entirely on high-resolution mass analyzers, whose precision and resolving power reliably distinguish complementary ion clusters from background signals. Compared to conventional MS3 quantification, a key limitation of TMTproC is its reduced multiplexing capacity: in most routine applications, only 9 of the 18 available TMTpro tags can be resolved^28^. This limitation, though, is not fixed; super-resolution techniques now enable the use of shorter transients combined with post-acquisition processing to achieve effective resolving powers of ≥500K, allowing extension to a 12-plex TMTproC setup without ultra-long transient acquisition^29^.

While cutting-edge techniques like super-resolution continue to expand the multiplexing capabilities of high-end instrumentation, it remains equally important to facilitate broader adoption on accessible and affordable mass spectrometers. Therefore, we explored the feasibility of adapting TMTproC methods to lower-resolution analyzers, specifically a quadrupole-ion trap instrument, to increase their utility in discovery-based proteomics.

Here, we introduce iTMTproC—an adaptation of TMTproC for quadrupole-ion trap instrumentation—enabling nine-channel multiplexed peptide quantification on a cost-effective, low-resolution platform. We benchmark iTMTproC against established multiplexed quantification methods, demonstrating comparable accuracy and sensitivity to MultiNotch MS3 workflows on Orbitrap Lumos™. Although data quality and sensitivity cannot match specialized multiplexing approaches like RTS-MS3 and RTS-TMTproC^24, 28^, our development represents a critical step toward democratizing multiplexed shotgun proteomics in laboratories lacking access to high-end mass spectrometry.

## Results and Discussion

### Optimizing Complement Reporter Ion Quantification in Ion Traps

To establish robust parameters for iTMTproC on quadrupole-ion trap instruments, we adapted previously described TMTproC methods, originally developed for quadrupole-Orbitrap™ analyzers^27^. Adjustments were necessary to accommodate the lower resolution characteristic of ion trap MS1 and MS2 scans. Additionally, we refined dynamic exclusion settings based on parameters optimized for label-free ion-trap experiments^30^. We evaluated peptide identification and quantification performance using a defined yeast peptide standard labeled with TMTpro at ratios of 0:1:5:10:1:10:5:1:0 across the nine complementary ion channels (Figure 2a). A significant challenge for implementing TMTproC in ion traps is the comparatively low mass accuracy during MS1 scans. This limited accuracy complicates quadrupole isolation, particularly in distinguishing closely spaced isotopic peaks (e.g., M0 and M+1). Consequently, multiple peaks can be inadvertently co-isolated, undermining accurate selection of the intended peak which is critical for reliable quantification in TMTproC workflows (Figure 2b). To address this challenge, we introduced a filtering step targeting doubly charged (2+) peptides. Specifically, we limited the quadrupole isolation window to ±0.2 Th around the theoretical M0 and M+1 peptide m/z values, significantly improving quantification accuracy (Figure 2c). In principle, triply charged (3+) precursor peptides could also be quantified, though this would require an even narrower quadrupole isolation window around theoretical M0 and M+1 peaks. However, minor calibration drifts of the quadrupole introduced noticeable quantification distortion for these peptides. To ensure robustness and reproducibility, we excluded 3+ precursor peptides from subsequent analyses in this study. This filtering step resulted in a 19% reduction in unique quantifiable peptides (from 7,359 to 5,995 peptides). Despite this decrease, we demonstrate subsequently that the overall number of quantified peptides and proteins remains highly competitive, confirming the effectiveness and practicality of this optimized approach.

**Figure 2.**
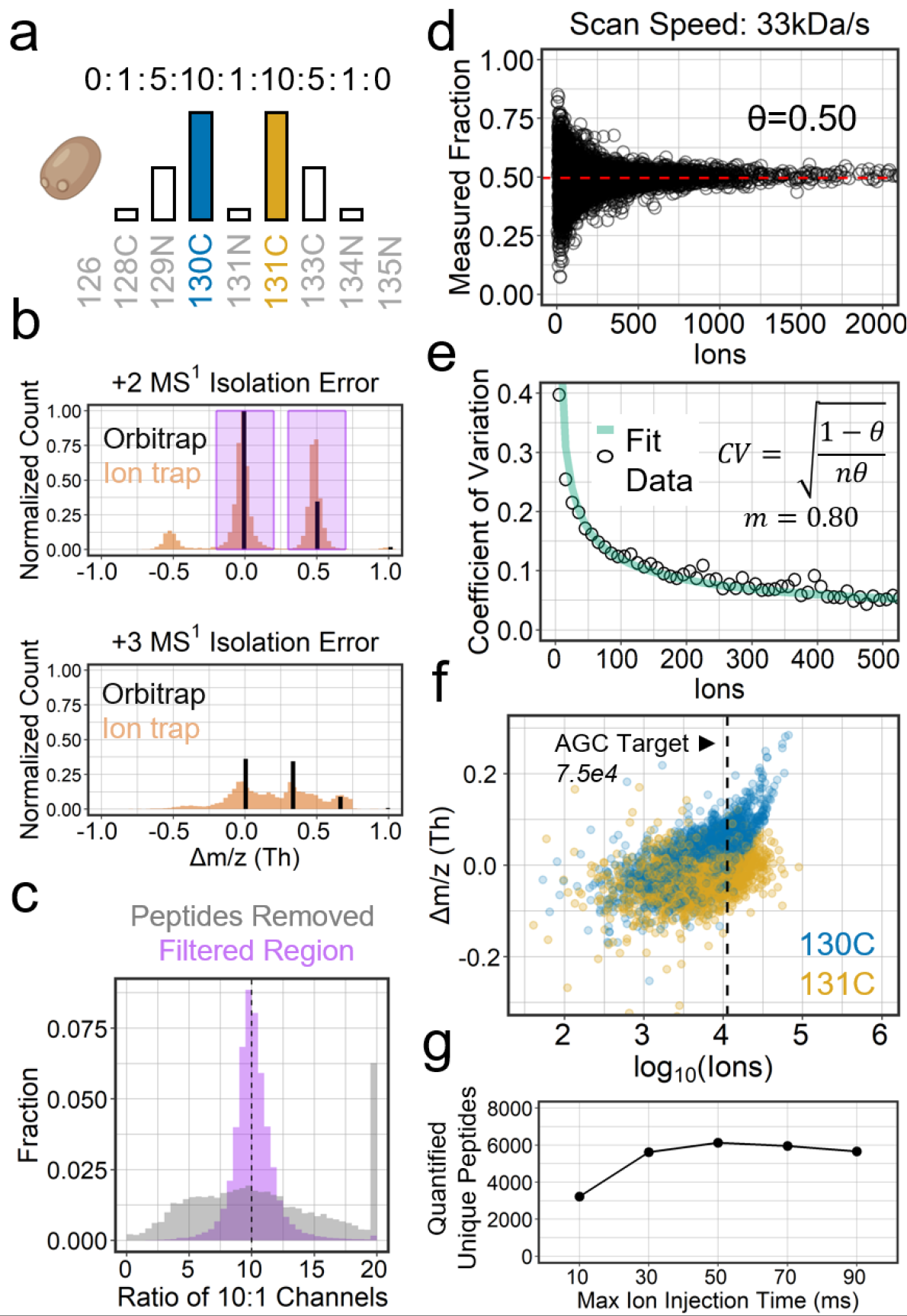
Method optimizations for complement reporter ion quantification in the ion trap. **(a)** Yeast standard. Yeast peptides were labeled with TMTpro™ in defined ratios of 0:1:5:10:1:10:5:1:0 across nine channels. This standard was used for all optimization experiments shown in this figure. **(b)** Histogram of Δm/z MS1 isolation errors for 2+ and 3+ peptides. The comparatively low-resolution MS1 scan in the ion trap leads to broader isolation errors when centering the isolation window for the subsequent MS2 scan. To address this, MS2 spectra were filtered to exclude cases where isolation was not centered on the M0 or M+1 precursor (i.e., peaks outside the highlighted purple region). Quantification of 3+ precursors is particularly challenging due to the narrow spacing between isotopes; therefore, all subsequent analyses were restricted to 2+ peptides. Frequency histograms are max-normalized and scaled by the relative charge state contributions. **(c)** Precision improvement achieved by filtering based on isolation m/z error. The purple histogram shows the distribution of peptide ratios retained after applying the isolation m/z filter (see panel b), with values tightly centered around the expected 10:1 ratio, indicating high measurement precision. In contrast, the grey histogram represents the excluded peptide data, which displays a broad distribution and high quantification error. **(d)** Relationship between observed peptide ratios and ion counts. Using the 10:10 TMTpro channels, the measured peptide fraction was plotted against the summed ion counts. The red dashed line indicates the median measured fraction (θ), defined as TMTpro130C / (TMTpro130C + TMTpro131C). The overall dataset median is 0.50, consistent with the expected 1:1 ratio. As more ions are sampled, measured ratios converge toward this expected value, reflecting improved statistical precision. **(e)** Coefficients of variation (CVs) at binned ion counts were used to fit a binomial model (equation shown), yielding a conversion factor of 0.80 (m) for translating ion trap charges into pseudo-counts (n). This factor remains approximately constant across different ion trap resolutions (see Figure S1). The conversion facilitates statistical analysis and helps define signal thresholds based on desired measurement precision. **(f)** Effect of ion trap signal intensity on channel error. Channel error is plotted against log_10_-transformed ion counts for the two most abundant complement reporter ions in the yeast standard (130C and 131C). To assess the impact of signal intensity, we intentionally overfilled the ion trap using an automatic gain control (AGC) target of 1.2e5. When ion counts exceed ~10,000 (1e4), systematic m/z shifts are observed, likely due to space charging. Based on this, we selected an AGC target of 7.5e4 for all subsequent experiments. The dotted line indicates the expected resulting number of complement ions, a regime that minimizes measurement deviation and supports reliable quantification. **(g)** Ion injection time optimization. With the AGC target fixed at 7.5e4 and scan speed set to 33 kDa/s, we varied the ion injection time (IIT) to determine its effect on peptide quantification. An IIT of 50 ms yielded the highest number of quantified unique peptides.

Next, we established the relationship between raw ion signals (denormalized by ion injection time) and pseudo-counts derived from a binomial statistical model. Defining this relationship is essential for setting appropriate signal thresholds corresponding to desired measurement precision^31^. To determine the conversion factor between ions and pseudo-counts, we analyzed peptide ratios from the equally mixed 10:10 TMTpro channels in our yeast standard, plotting these ratios against their summed ion counts (Figure 2a, 2d). With fewer ions sampled, the measured ratios showed high variability due to statistical fluctuations. As ion sampling increased, these ratios converged toward the expected 1:1 ratio, consistent with predictions from the binomial model. From the relationship between the expected coefficient of variation (CV) and pseudo-counts, we derived a conversion factor of approximately 0.8 pseudo-counts per ion. We confirmed the consistency of this conversion factor across various ion trap scan speeds, ranging from 66 to 2 kDa/s (Rapid, Normal, Enhanced, and Zoom; Figure S1). This finding contrasts with Orbitrap™-based methods, where higher analyzer resolution typically results in smaller conversion factors^31-33^. Based on this analysis, we established a minimum pseudo-count threshold of 450 in the nine resolvable TMTproC channels, corresponding to an expected coefficient of variation (CV) of ~10% for a 1:1 ratio across all channels.

We then optimized two additional critical parameters: the automatic gain control (AGC) and ion injection time (IIT) specifically for iTMTproC. Space charging poses a significant limitation in ion trap mass analyzers, similar to the Orbitrap™^32, 34^. If too many ions accumulate simultaneously, ion-ion interactions distort mass accuracy. Unlike Orbitrap™ analyzers, however, the lower resolution spectra obtained from ion traps cannot be effectively recalibrated post-acquisition to correct such distortions. To empirically determine the maximum ion capacity of the ion trap, we intentionally overloaded it using a high AGC target of 1.2e5. We then examined the mass accuracy of complement reporter ions as a function of the total ion intensity within their clusters (Figure 2f). This analysis revealed an approximate threshold above which mass accuracy began to deteriorate. Considering the typical complement ion generation efficiency of roughly 15%^27^, this threshold corresponded to an optimal AGC target of approximately 7.5e4. Consequently, we adopted this AGC setting for all subsequent analyses.

Lastly, we optimized ion injection time by varying IIT values systematically from 10 to 90 ms (Figure 2g). We assessed each IIT setting based on the number of uniquely quantified peptides meeting our previously established quality criteria (pseudo-count >450 and ±0.2 Th isolation window filter). An IIT of 50 ms yielded the highest number of quantified peptides. Taken together, these experiments established the final AGC (7.5e4) and IIT (50 ms) parameters for iTMTproC, enabling robust and accurate quantification on quadrupole-ion trap instrumentation. Detailed experimental procedures outlining these optimized settings are provided in the methods section and were applied throughout the remainder of this study.

### Evaluating Stellar iTMTproc with an interference standard against Orbitrap Lumos™ MultiNotch MS3

Next, we evaluated iTMTproC with a standard that combines two samples (yeast and human) with differing mixing ratios^19, 20^ (Figure 3a). We combined one part yeast versus ten parts human to simulate interference for biologically dynamic proteins, which tend to be lower abundant^26, 35^. We have chosen to compare those results with MultiNotch MS3 on the Orbitrap Lumos™ due to its proven track record to generate highly valuable data that has led to numerous biological insights^36-41^. First, we varied ion trap scan speeds from 66 (Rapid) to 2 kDa/s (Zoom) and determined how this would affect the observed interference (Figure S2). Higher resolution helps to distinguish real signal from background noise. However, the higher resolution in the ion trap comes at the cost of dramatically slower scan speeds (Figure S2e). Based on our previous finding that our desired IIT is 50ms, we chose 33 kDa/s (Normal) for iTMTproC analysis, which takes approximately 36 milliseconds to scan an MS2 spectrum with 1200 m/z range (Figure S2e). Importantly, ion injection time and ion trap m/z analysis are parallelized on the quadrupole-ion trap instruments and the 33 kDa/s scanning resolution therefore does not significantly slow down iTMTproC analysis.

**Figure 3.**
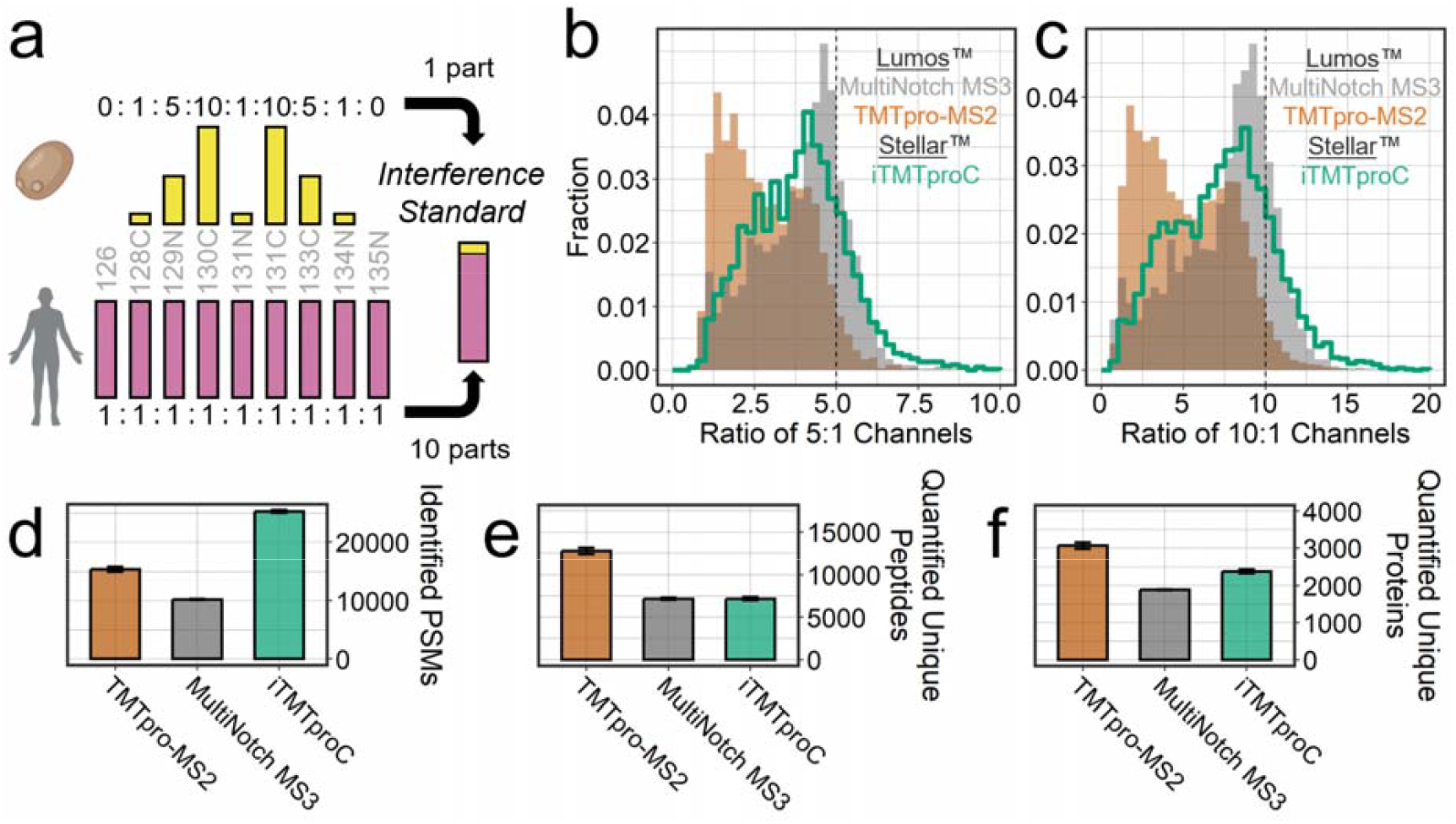
Multiplexed iTMTproC shotgun proteomics on Stellar™ yields results comparable to MultiNotch MS3 on the Orbitrap Lumos™. **(a)** Human-yeast interference standard. Yeast peptides were labeled with TMTpro™ in defined ratios of 0:1:5:10:1:10:5:1:0 and combined with HeLa peptides labeled 1:1:1:1:1:1:1:1:1 across the same channels. To simulate the interference encountered with dynamic, typically low-abundance proteins, the samples were mixed at a 1:10 yeast-to-human ratio. **(b-c)** Ratio distortion comparison of iTMTproC on Stellar™ vs MultiNotch MS3 on the Orbitrap Lumos™. Histograms of measured ratios for the 5:1 (b) and 10:1 (c) yeast channels. Ratios closer to the expected values indicate reduced interference and improved quantification accuracy. While MultiNotch MS3 on the Orbitrap Lumos™ shows slightly higher accuracy, both methods substantially outperform TMTpro-MS2– based quantification on the Orbitrap Lumos™. **(d)** Number of peptide-spectrum matches (PSMs). iTMTproC yields the highest number of PSMs, driven by its fast scan speed, the high sensitivity of the ion trap, and repeated targeting of the same isotopic envelope. Error bars represent the standard error of the mean (n = 3). **(e)** Number of quantified unique peptides. TMTpro-MS2 yields the highest number of unique quantified peptides; however, these measurements are often severely distorted due to interference. The number of unique quantified peptides is nearly identical between Stellar™ iTMTproC and Orbitrap Lumos™ MultiNotch MS3. Although iTMTproC shows a lower conversion rate from PSMs to unique peptides—due to stringent filtering and redundancy—this is compensated by its higher total PSM count, resulting in comparable peptide-level coverage. **(f)** Number of quantified proteins. TMTpro-MS2 on the Orbitrap quantifies the highest number of proteins, though these measurements suffer from severe ratio distortion. On average, Stellar™ iTMTproC quantifies 26% more proteins than Orbitrap Lumos™ MultiNotch MS3 in a single shot analysis.

Next, we assessed quantification accuracy by examining measured ratio distributions for yeast peptides in the 5:1 and 10:1 channels, which serve as indicators of interference levels (Figure 3b, c). While MultiNotch MS3 on the Orbitrap Lumos™ exhibited slightly less ratio distortion, Stellar™ iTMTproC minimizes interference, producing distributions that closely align with MS3 data. Significantly, both methods represent a substantial improvement over TMTpro-MS2 reporter ion quantification in the Orbitrap™, where low-m/z reporter ions are more susceptible to co-isolation interference.

We also evaluated method sensitivity and found that iTMTproC on Stellar™ quantifies about the same number of unique peptides as Orbitrap Lumos™ MS3 while quantifying 30% more proteins (Figure 3d-f). This is likely due to iTMTproC’s capability to quantify lower abundant 2+ peptides, while MultiNotch MS3 can quantify 3+ peptides, which will add new unique peptides but tend to come from the highest abundant proteins that were already quantified. It is important to note that this difference in protein identifications is most pronounced in single-shot analyses; in applications where prefractionation is used to distribute peptides across multiple fractions, the difference becomes less pronounced but still noticeable, as shown in a later section.

Comparing methods between different instruments, associated HPLCs, and laboratories has the inherent caveat that it is unclear how much of the observed differences are due to the actual methods, versus instrument or HPLC calibration, instrument performance, and setups. To minimize this possibility for biases, we evaluated iTMTproC, TMTpro-MS2, and MultiNotch MS3 (Non-RTS) on a single HPLC (Vanquish Neo™) and a single mass spectrometer (Ascend™) (Figure S3). The Ascend™ quadrupole and low-pressure ion trap used for m/z analysis are identical to the standalone quadrupole-ion trap (Stellar™), though the Ascend™ incorporates a larger high-pressure trap to support higher space charge capacity for techniques such as electron transfer dissociation. Compared to the Orbitrap Lumos™, the Ascend™ quadrupole is a significant advancement with higher ion transfer efficiency and a more ideal, rectangular transmission profile^42^. The increased selectivity and sensitivity are expected to result in data from MS3 and TMT-MS2 that would outperform data acquired with these methods on the Orbitrap Lumos™. Nevertheless, comparing iTMTproC, MultiNotch MS3, and TMTpro-MS2 on a single instrument mirrored the results of the comparison between Stellar™ and Orbitrap Lumos™ — iTMTproC exhibited higher sensitivity than MultiNotch MS3 but slightly lower accuracy, while still greatly improving data quality over TMTpro-MS2 (Figure S3). Overall, we conclude that iTMTproC is able to obtain accurate multiplexed proteomics data with slightly more ratio distortion than MultiNotch MS3. However, this is compensated by a slightly larger number of quantified proteins.

### Evaluating iTMTproC and MultiNotch MS3 in a Complex Biological Sample

To evaluate the practical applicability of iTMTproC beyond controlled standards, we tested its performance using complex biological samples. Specifically, we assessed protein dynamics during early embryonic development of the frog *Xenopus laevis*, tracking changes from fertilized eggs to swimming tadpoles (Figure 4). At each developmental time point, we collected four embryos, prepared protein samples following established protocols, and performed pre-fractionation via medium pH reverse-phase chromatography^43, 44^. We then analyzed 24 fractions using both iTMTproC and MultiNotch MS3 on the Ascend™ instrument. Both methods quantified similar numbers of unique peptides; however, iTMTproC identified 8,971 proteins compared to 8,531 proteins quantified by MultiNotch MS3—approximately a 5% increase in proteome coverage, highlighting iTMTproC’s enhanced sensitivity (Figure 4b). The proteins identified by both methods showed substantial overlap (Figure 4c). To further evaluate proteome coverage, we compared quantified proteins against a reference dataset of absolute protein abundances measured previously in *Xenopus laevis* eggs^45^. Protein abundance distributions were nearly identical across methods (Figure 4d), confirming similar dynamic range coverage. Notably, the total number of proteins quantified by either approach surpasses previously published results from our lab using a similar developmental time-series analyzed with MultiNotch MS3 (TMT-10 plex) on an Orbitrap Lumos™, which quantified 7,827 proteins after reanalysis with updated gene models^46^. These findings demonstrate that iTMTproC delivers data quality and proteome coverage comparable to what was achievable with then state-of-the-art multiplexed shotgun proteomics methods just a few years ago.

**Figure 4.**
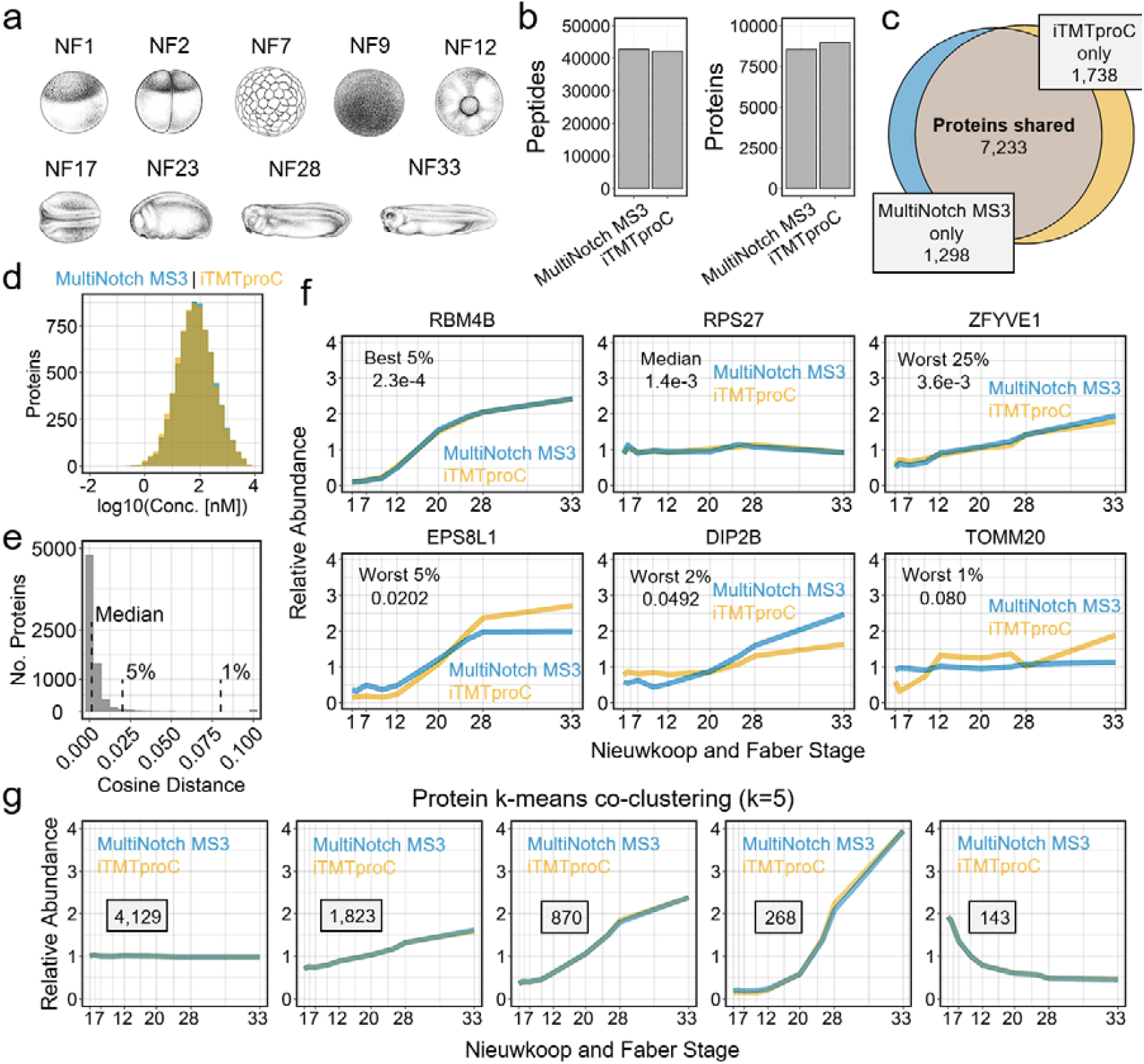
Comparing iTMTproC with MultiNotch MS3 to analyze proteomic changes across developing frog embryos. **(a)** Developmental stages of the frog *Xenopus laevis*. Embryos were collected at defined Nieuwkoop and Faber stages and labeled with TMTpro™ reagents for quantitative proteomic analysis using both iTMTproC and MultiNotch MS3^49^. Xenopus illustrations from Natalya Zahn^50^. **(b)** Quantified unique peptides and proteins from 24 fractions. MultiNotch MS3 quantified slightly more unique peptides, while iTMTproC quantified approximately 5% more proteins. **(c)** Venn diagram of quantified proteins. Most proteins are quantified by both methods, with a slightly larger number uniquely quantified by iTMTproC. **(d)** Distribution of absolute protein concentrations. Both iTMTproC and MultiNotch MS3 capture a similar dynamic range of protein abundances^45^. **(e)** Cosine distance histogram between iTMTproC and MultiNotch MS3 measurements. Histogram of cosine distances for proteins quantified by both methods. Vertical lines indicate the median (distance = 1.4e-3), 95th, and 99th percentiles. The distribution shows that most proteins exhibit highly similar expression profiles, independent of the quantification method. **(f)** Representative examples of protein quantification. Shown are proteins at the 5th percentile (RBM4B), median (RPS27), 75th percentile (ZFYVE1), 95th percentile (EPS8L1), 98th percentile (DIP2B), and 99th percentile (TOMM20) based on cosine distance between the two methods. Even proteins in the most distant 2% exhibit similar expression trends. Only the most divergent 1% show substantial disagreement. The x-axis is scaled by hours post-fertilization. **(g)** Protein k-means co-clustering (k = 5). Co-clustering of proteins across paired developmental time points reveals nearly identical expression dynamics between iTMTproC and MultiNotch MS3, highlighting the high concordance and comparable data quality between the two methods.

While the number of quantified proteins is promising, even more critical is the agreement between protein dynamics measured by different quantification methods. To assess this quantitatively, we calculated cosine distances across the nine developmental time points for all proteins quantified by both iTMTproC and MultiNotch MS3 (Figure 4e). The resulting distribution of cosine distances showed a median value of 1.4e-3, indicating high overall similarity between temporal protein profiles obtained from the two approaches. However, interpreting cosine distances biologically can be challenging. To illustrate the biological meaning of these metrics more intuitively, we selected proteins representing the 5th, 50th, 75th, 95th, and 99th percentiles of agreement (with lower percentiles reflecting closer agreement) and plotted their developmental dynamics (Figure 4f). Encouragingly, protein dynamics remained essentially indistinguishable up to the 75th percentile of proteins. Even proteins at the 95th (EPS8L1) and 98th percentiles (DIP2B) exhibited similar overall dynamic trends. Only at the extreme (99th percentile, TOMM20) did substantial disagreement between the two methods become apparent. Given that both datasets were stringently controlled at a 1% false discovery rate (FDR) at both the peptide and protein levels, such discrepancies at the extreme percentile are expected.

The consistency of these measurements between iTMTproC and MultiNotch MS3 was further supported by global trends in protein expression. K-means clustering (k=5) of protein dynamics revealed highly similar developmental expression profiles across all clusters between the two methods (Figure 4g). Collectively, these analyses emphasize that, in practical biological applications, the systems-level data generated by iTMTproC on quadrupole-ion trap instrumentation and MultiNotch MS3 are essentially equivalent.

## Conclusions

In this study, we demonstrated that complement reporter ion quantification can be successfully adapted to quadrupole–ion trap instrumentation for multiplexed shotgun proteomics (iTMTproC). Benchmarking with defined standards and complex biological samples revealed that iTMTproC achieves quantification accuracy and interference reduction comparable to established MultiNotch MS3 methods on high-end Orbitrap™ instruments. Although iTMTproC currently exhibits higher ratio compression and lower sensitivity compared to the latest specialized multiplexing approaches^24, 28^, it importantly enables multiplexed quantitative analysis on cost-effective quadrupole–ion trap platforms. This advance significantly expands accessibility and helps democratize advanced quantitative proteomics. Until now, quadrupole–ion trap instruments such as the Stellar™ have primarily been employed for targeted analyses. In this context, iTMTproC represents an attractive option for increasing sample multiplexing without compromising assay throughput, as often encountered with slower MS3-based methods^47^. Our findings also highlight previously underappreciated multiplexing capabilities of ion-trap analyzers, which traditionally have been confined to qualitative or targeted proteomic applications^11, 48^. In future work, it will be particularly interesting to investigate whether the multiplexing strategy demonstrated here can be effectively applied to multiplexed targeted proteomics workflows.

In this study, we implemented iTMTproC by isolating ions using the quadrupole. However, ion traps themselves can, in principle, also achieve effective ion isolation. In configurations with adjacent higher-energy collision dissociation (HCD) cells, where ions are injected simultaneously into the trap, space-charging effects can become particularly challenging—especially for the narrow isolation windows required by iTMTproC—reducing isolation effectiveness and quantification accuracy. Nevertheless, this limitation could potentially be overcome in instruments designed for continuous ion transfer into the trap. In future work, we plan to evaluate whether robust iTMTproC quantification can be achieved on ion-trap-only instruments. Demonstrating this capability would further reduce instrument costs and has the potential to significantly expand access for quantitative proteomics in non-specialized laboratories.

**Figure S1.**
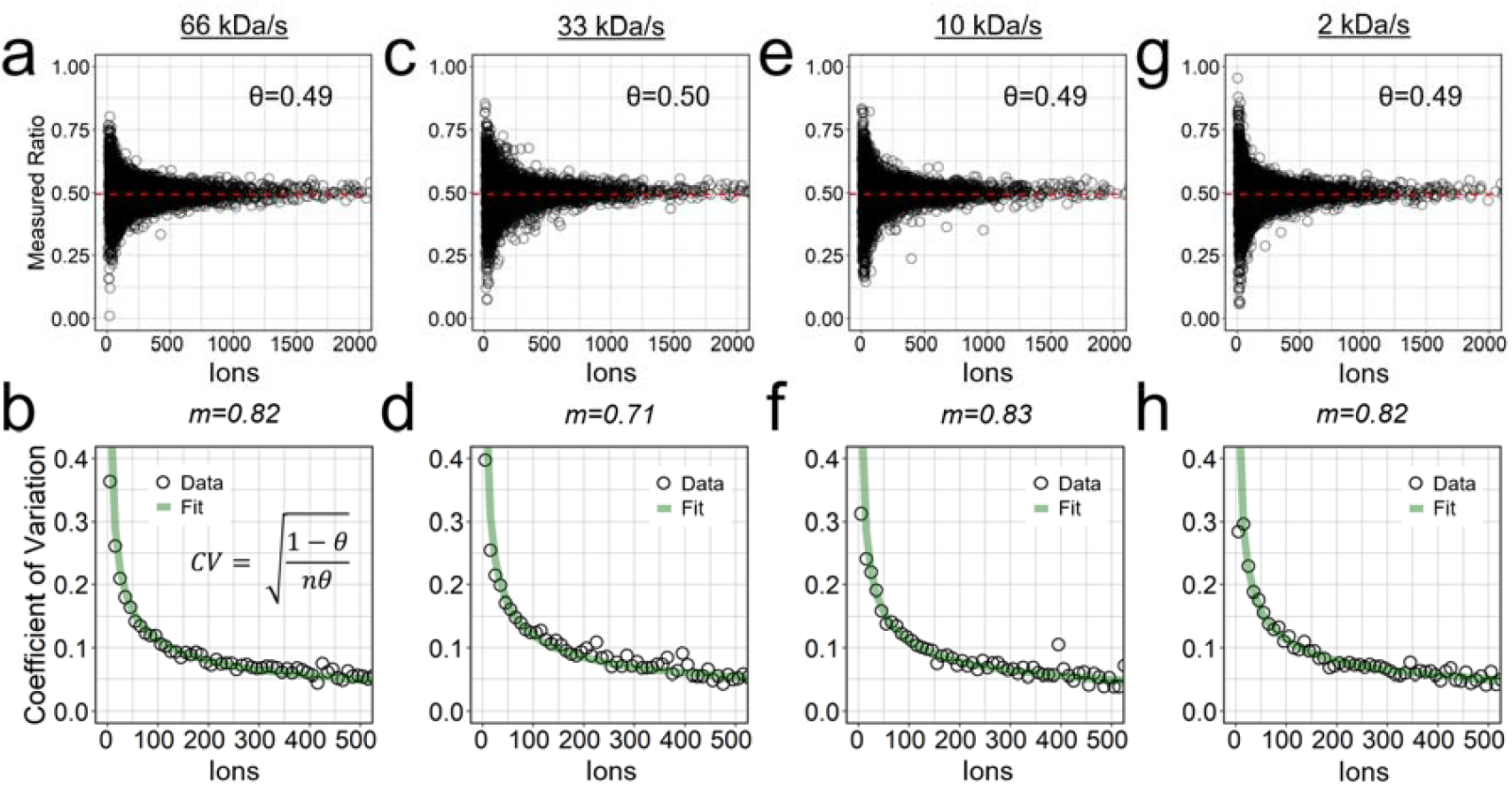
Scan speed does not affect the conversion factor between charges and pseudo-counts in an ion trap. Panels (a, c, e, g) show measured peptide ratios plotted against summed charges at different scan speeds (66, 33, 10, and 2 kDa/s, respectively). The expected ratio (θ) remains consistent across all conditions. Panels (b, d, f, h) display the coefficients of variation (CVs) as a function of charges for each corresponding scan speed. The conversion factor, derived from binomial modeling, remains stable across scan speeds, confirming that—unlike in the Orbitrap™—this factor does not depend on resolution in an ion trap. All data were acquired in a single run, testing all four scan speeds in parallel.

**Figure S2.**
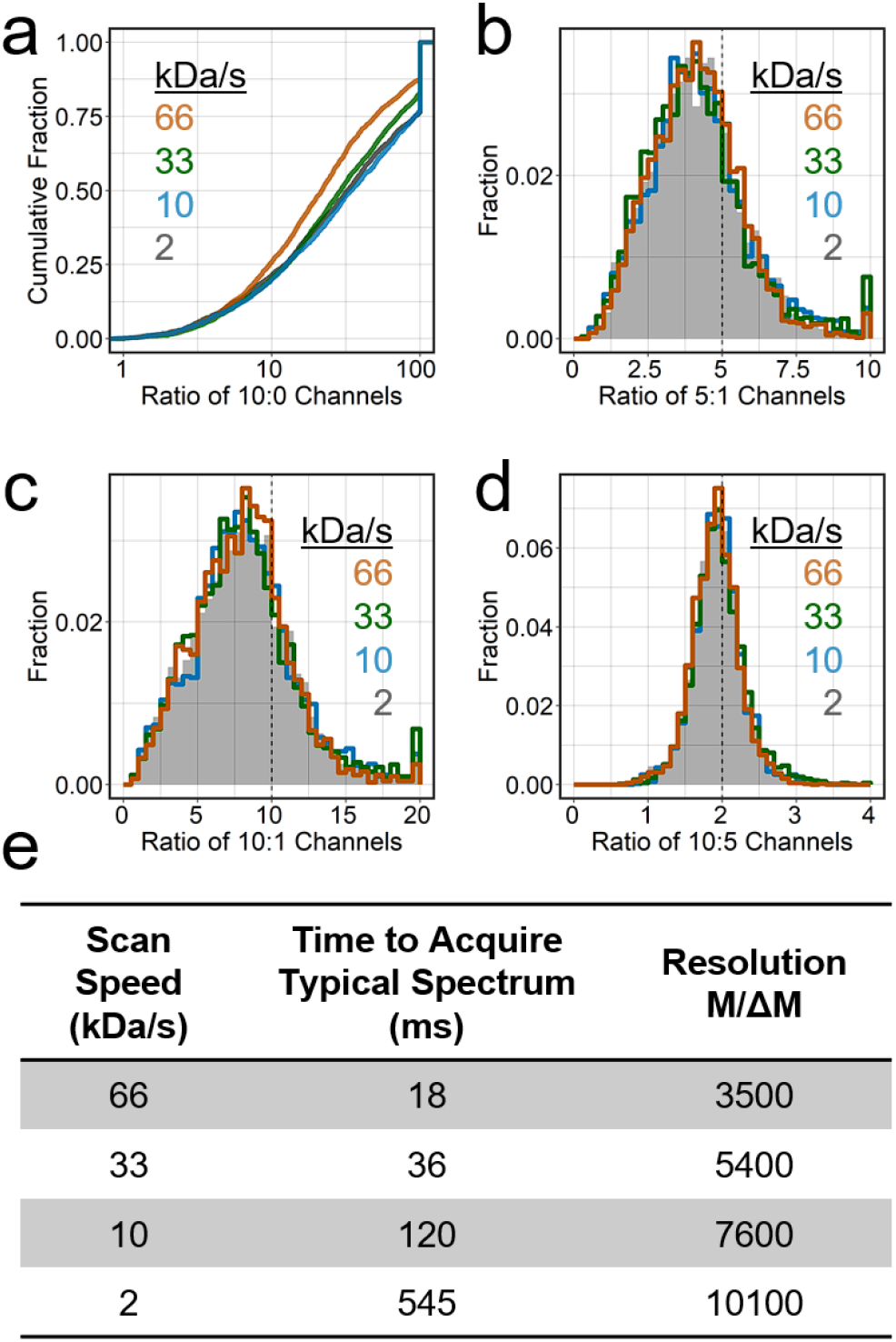
Evaluation of ion trap scan speeds for iTMTproC quantification using the human– yeast interference standard. Four ion trap scan speeds (66, 33, 10, and 2 kDa/s) were tested in a single run to determine the fastest speed that maximizes peptide identifications while maintaining data quality. (a) Cumulative distribution of measured 10:0 channel ratios. Lower cumulative fractions indicate reduced interference. (b–d) Histograms of measured ratios for the interference standard: 5:1 (b), 10:1 (c), and 10:5 (d). Results show that scan speed has minimal impact on resolving complement peaks across all speeds tested. However, 66 kDa/s was found to be sensitive to calibration drift and was deemed impractical for routine use. (e) Summary table of manufacturer-reported scan speed specifications. For each scan speed, the corresponding time to acquire a typical 1200 m/z spectrum and the reported resolution (M/ΔM) are listed. Based on these findings, 33 kDa/s was selected as the standard scan speed for iTMTproC due to its optimal balance of speed, resolution, and robustness.

**Figure S3.**
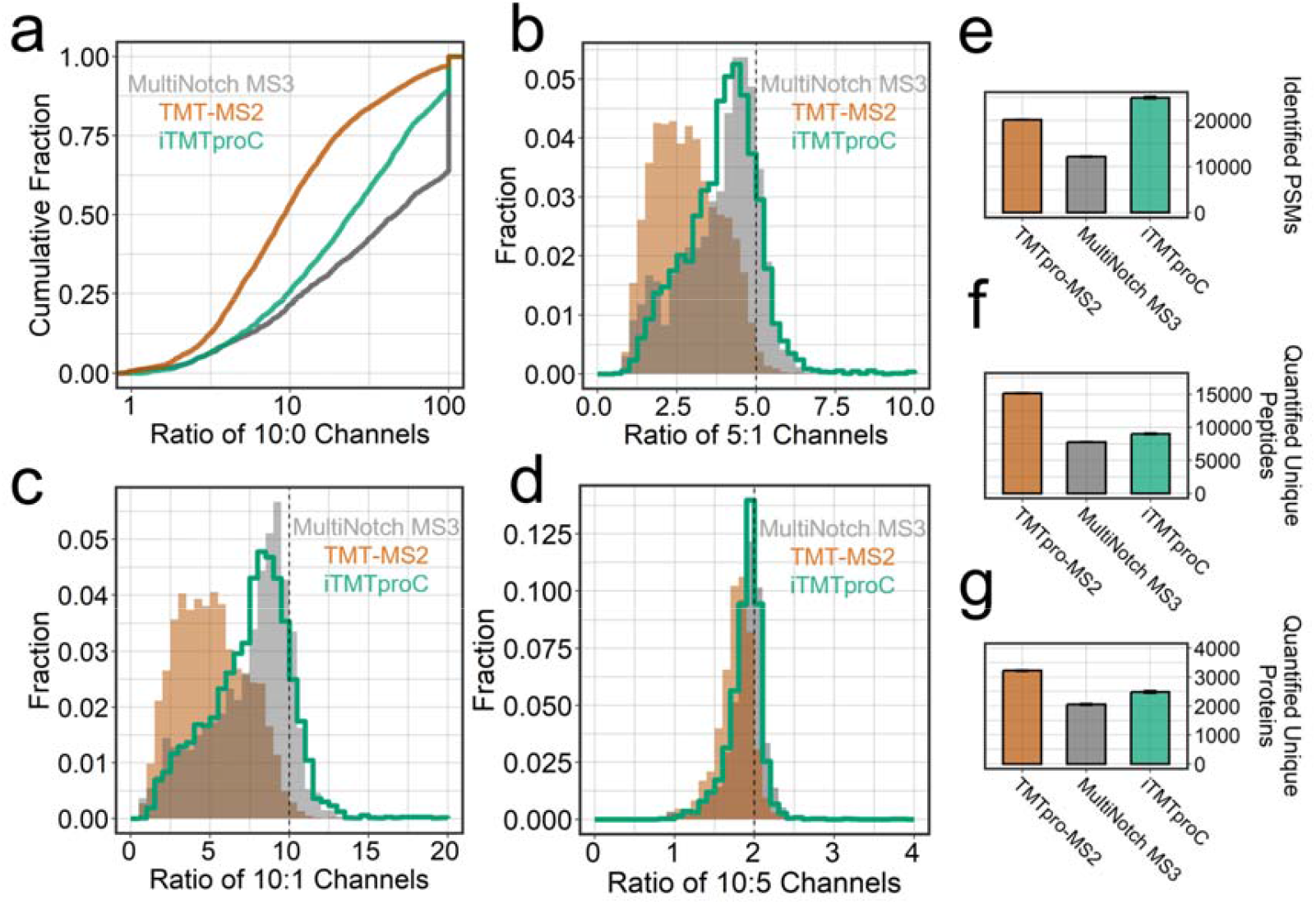
Comparison of iTMTproC with MultiNotch MS3 and TMTpro-MS2 on the Ascend™. (a) Cumulative distribution of the measured 10:0 channel ratios. Since the true 10:0 ratio is infinite, a lower cumulative fraction less than 100 suggests reduced interference. TMTpro-MS2 shows the most distortion, and ion trap TMTproC (iTMTproC) outperforms TMTpro-MS2 but is still behind MultiNotch MS3. Data for iTMTproC, TMTpro-MS2, and MultiNotch MS3 was acquired on an Ascend™, ensuring a fair comparison of all methods using a consistent gradient and an instrument capable of accurately representing all three approaches. The Ascend™ has a brighter beam and more parallelizable injection time than a Fusion™ series MS. However, for FTMS2 methods, it is reasonable to compare this to data quality achieved on an Orbitrap Lumos™. (b-d) Histograms of measured ratios for 5:1 (b), 10:1 (c), and 10:5 (d) channels. Ratios closer to the expected values indicate less interference and improved accuracy. Further illustrating the trend observed in (b), iTMTproC exhibits comparable interference reduction as MultiNotch MS3 on the same instrument. (e–g) Peptide and protein identification. Like analyses on independent instruments, iTMTproC yields the highest number of PSMs and quantifies more unique proteins than MultiNotch MS3. As an intermediate approach, iTMTproC offers higher sensitivity than MultiNotch MS3 but slightly lower data quality, while greatly improving data quality over TMTpro-MS2 with a reduction in sensitivity. Error bars are shown and represent the standard error of the mean (n = 3).

## Experimental Procedures

### Proteomics Sample Preparation

Proteomics samples were prepared as previously described^27, 44, 46^. In brief, following cell lysis, samples are reduced with 5mM dithiothreitol (DTT) for 20 minutes at 60°C and alkylated with 20 mM *N*-ethylmaleimide for 20 minutes at room temperature. 5mM DTT was added to quench the excessive alkylating reagents. Proteins were purified using one of two methods, depending on the sample: (1) For our HeLa and yeast samples, proteins were purified by methanol-chloroform precipitation^51^, and the resulting dried pellet was resuspended in 10 mM *N*-(2-hydroxyethyl)piperazine-*N*′-3-propanesulfonic acid (EPPS; pH 8.5) with 6□M guanidine hydrochloride. Samples were heated at 60□°C for 15□min, and the protein mixture was diluted threefold with 10□mM EPPS (pH 8.5). (2) For our *Xenopus laevis* samples, proteins were precipitated following reduction and alkylation using the SP3 method, as previously described^52^. After binding and washing the bead-bound protein, the protein-containing beads were resuspended in 2□M guanidine hydrochloride. Following protein precipitation, the protein mixture was digested with 20 ng/µL LysC (Wako) overnight at room temperature. Samples were further diluted fourfold with 10 EPPS (pH 8.5) and digested with an additional 20 ng/µL LysC and 10 ng/µL sequencing-grade trypsin (Promega) at 37°C for 16h. After digestion, the peptides were cleared by ultracentrifugation at 100,000g for 1h at 4°C, and the supernatant was vacuum dried. For TMTpro labeling, TMTpro tags were added at a ratio of 5□μg of TMTpro:1□μg of peptide, mixed, and incubated at room temperature for 2□h. The reaction was then quenched by addition of 5□μL of 5% hydroxylamine at room temperature for 30□min. The resulting mixture was vacuum dried, desalted using homemade stage tips with C18 material (Empore), and resuspended in 1% formic acid to 1 µg/µL before LC–MS analysis.

For the *Xenopus* development time course, prefractionation was utilized to detect a larger number of peptides^43^. Specifically, before LC–MS analysis, the dried peptides were resuspended in 10□mM ammonium bicarbonate (pH 8) with 5% acetonitrile to a peptide concentration of 1□μg/μL. The dissolved peptides were separated into 96 fractions using medium pH reverse-phase separation (Zorbax 300Extend C18, 4.6□×□250□mm column) on a 1260 Infinity II LC system (Agilent), as described previously^43^. Each resulting 96-well plate was combined into 24 fractions, and each fraction was desalted and resuspended for LC–MS analysis, as described above.

### HeLa-yeast interference standard

HeLa S3 cells were grown on 10 cm tissue culture plates to 80% confluency, and *S. cerevisiae* S288C was grown to an OD of 0.4 in YPD as a suspension culture. HeLa cells were pelleted and lysed by sonication in 100 mM HEPES buffer (pH 7.2) with 2% SDS and Roche protease inhibitor. Yeast cells were lysed by cryomilling and later resuspended in 50 mM HEPES (pH 7.2) with 4% SDS and 1 mM DTT. Samples were further prepared as described above. For TMTpro labeling, yeast lysate was labelled at 0:1:5:10:1:10:5:1:0 ratios using the nine TMTproC-compatible channels, and the HeLa lysate was labelled as 1:1 using the same channels. Prior to desalting, the TMT-labelled yeast and HeLa lysate are mixed at a 10 HeLa:1 yeast ratio.

### Collection of developing Xenopus laevis embryos

Mature *X. laevis* females and males were purchased from Xenopus1 and maintained by Laboratory Animal Resources at Princeton University. All animal procedures are approved under Institutional Animal Care and Use Committee protocol 2070. Unfertilized eggs and male testes were collected following standard laboratory procedures previously described^53^. For testes collection, *X. laevis* males are euthanized in 0.1% (w/v) tricaine methanesulfonate (MS-222; Syndel’s Syncaine) and then sacrificed by pithing. The testes are isolated and stored at 4 °C in oocyte culture medium that was exchanged daily for up to 1 week [1 L: 13.7 g Leibovitz’s L-15 Medium Powder (Thermo Fisher Scientific; #41300039), 8.3 ml penicillin–streptomycin (Thermo Fisher Scientific; #15140122), 0.67 g bovine serum albumin]. For egg collection, female frogs were injected with 500 U of human chorionic gonadotropin (CG10; Sigma) and kept at 16 °C in Marc’s modified Ringer’s solution for 16 h before collection (1X MMR: 5 mM HEPES (pH 7.8), 0.1 mM EDTA, 100 mM NaCl, 2 mM KCl, 1 mM MgCl2, and 2 mM CaCl2). For in vitro fertilization, female eggs collected in 1X MMR buffer are cleaned, and preactivated eggs are removed. Half of one male testis was used per 500 eggs by crushing in 1X MMR buffer with a sterile pestle and then mixing with the unfertilized eggs. The mixture was incubated at 16 °C for 5 min, followed by mixing and an additional 5 min incubation. Fertilization was induced by flooding the eggs with 0.1X MMR. After 1 h at 16 °C, embryo jelly coats were removed by incubating with 2% cysteine in 0.1X MMR for 5 min, and the embryos were thoroughly washed with 0.1X MMR to remove residual cysteine. Embryos were grown and staged by Nieuwkoop and Faber^49^ nomenclature at 16 °C and then flash frozen at desired time points. Embryo lysis was performed as described previously^46^ and the subsequent sample preparation protocol is stated above.

### UHPLC Chromatography

All proteomics samples were analyzed on a Vanquish Neo™ UHPLC System or Proxeon 1200. Solvent A consisted of 2% DMSO and 0.125% formic acid in water, and solvent B consisted of 80% acetonitrile, 2% DMSO and 0.125% formic acid in water. Depending on the experiment, the UHPLC system was coupled to either an Ascend™ (Vanquish), Orbitrap Lumos™(Proxeon), or Stellar™(Vanquish) mass spectrometer. Notably, comparisons between Orbitrap Lumos™ and Stellar™ utilize entirely separate UHPLC systems at two distinct locations. On all instruments, peptides were separated on an Aurora Series emitter column (25□cm□×□75□μm inner diameter, 1.6-μm C18; IonOpticks) and were held at 60□°C using an in-house-built column oven while experiments with the Stellar utilized a Nanospray Flex™ ion source (Thermo Fisher).

All samples, were analyzed with the following 90-min gradient at a constant flow rate of 350□nL/min after thorough equilibration of the column to 0% B: 0–10% B in 5□min, 10–26.4% B for 70□min, 26.4–100% for 10□min and 100% for 5□min. For electrospray ionization, 2.6□kV was applied between 1□min and 83□min of the LC gradient. To avoid carryover of peptides, 2,2,2-trifluoroethanol was injected in a 30-min wash between each sample^54^. For fractionated samples, this wash was performed between every three fractions from the same original sample.

### Ion trap complement reporter ion quantification (iTMTproC) Method

iTMTproC optimization experiments were performed on the Ascend™. The instrument designation based on experiment is as follows: Optimization of iTMTproC parameters – Ascend™ (Figure 2/S1/S2), Interference Standard Comparison with MultiNotch MS3 – Stellar™ (Figure 3), *Xenopus* Developmental Proteomics – Ascend™ (Figure 4).

The mass spectrometer was set to analyze positively charged ions in a data-dependent MS2 mode, recording centroid data with the RF lens level at 60%. Full scans were taken with the ion trap at 33 kDa/s (Normal resolution) with an automatic gain control (AGC) target of 3e4 ions, maximum IIT of 30 ms, and scan range of 500 to 1200 *m/z* with wide quadrupole isolation enabled. Maximum cycle time between MS1 scans was set to 3 s.

Following the survey scan, the following filters were applied for triggering MS2 scans. Isolated masses were excluded for 15 s after triggering a mass tolerance window of ±0.5 m/z. Ions were analyzed if their *m/z* ratio was between 550 and 1050 to ensure visibility of the complementary ion clusters in a normal range MS2 scan. An intensity threshold was set for 5E3 ions.

The following settings were used for ion trap MS2 scans. AGC target was set to 7.5E4 charges, and the maximum IIT was 50 ms. The quadrupole was utilized for isolation with an isolation width of 0.4 Da, and ions were fragmented with collision induced dissociation (CID) at a normalized collision energy of 35% (10 ms activation time, 0.25 activation Q). The scan rate was set to 33 kDa/s (Normal resolution) on auto scan range mode with a default charge state of 2.

### MultiNotch MS3 Method

MultiNotch MS3 experiments were performed on the Orbitrap Lumos™ and Ascend™. The instrument designation based on experiment is as follows: Interference Standard Comparison with iTMTproC – Orbitrap Lumos™ (Figure 3), Interference Standard Comparison with iTMTproC – Ascend™ (Figure S2), *Xenopus* Developmental Proteomics – Ascend™ (Figure 4).

The mass spectrometer was set to analyze positively charged ions in a data-dependent MS3 mode, recording centroid data with the RF lens level at 60%. Full scans were taken with the Orbitrap™ at 120k resolution with an automatic gain control (AGC) target of 4E5 charges, maximum IIT of 50 ms, and scan range of 400 to 1600 m/z with wide quadrupole isolation enabled. Maximum cycle time between MS1 scans was set to 3 s.

Following the survey scan, the following filters were applied for triggering MS2 scans. Monoisotopic peak selection was enabled and set to isolate the most abundant peak in peptide mode. Isolated masses were excluded for 60 s after triggering with a mass tolerance window of ±10 ppm, while also excluding isotopes and different charge states of the isolated species. A charge state filter was set for 2-6+ charge states. An intensity threshold was set for 5E3 ions.

The following settings were used for ion trap MS2 scans. AGC target was set to 1E4 charges, and the maximum IIT was 35 ms. The quadrupole was utilized for isolation with an isolation width of 0.7 Da, and ions were fragmented with CID at a normalized collision energy of 35% (10 ms activation time, 0.25 activation Q). The scan rate was 33kDa/s (Normal resolution) in auto scan range mode.

For triggering MS3-scans for TMT-reporter ion-based quantification, the following filters were applied. Precursor ion exclusion was set to 5 (high) and 50 (low) m/z. Isobaric Tag Loss Exclusion was set to TMTpro, and 10 notches were used to isolate SPS-MS3-precursors. MS3 scans were acquired with a 0.7 m/z MS1 isolation window and 2 m/z MS2 isolation window. The Orbitrap™ detector was used with a resolution of 45k and an AGC target of 2E5, scanning over the range 100-500 m/z. The maximum ion injection time was 86 ms. Ions were fragmented in the higher-energy collision dissociation (HCD) cell at a normalized collision energy of 45%.

### TMT-MS2 Method

TMT-MS2 experiments were performed on the Orbitrap Lumos™ and Ascend™. The instrument designation based on experiment is as follows: Interference Standard Comparison with iTMTproC – Orbitrap Lumos™ (Figure 3), Interference Standard Comparison with iTMTproC – Ascend™ (Figure S2). The mass spectrometer was set to analyze positively charged ions in a data-dependent MS2 mode, recording centroid data with the RF lens level at 60%. Full scans were taken with the Orbitrap™ at 120k resolution with an automatic gain control (AGC) target of 4E5 charges, maximum IIT of 50 ms, and scan range of 350 to 1400 m/z with wide quadrupole isolation enabled. Maximum cycle time between MS1 scans was set to 3 s.

Following the survey scan, the following filters were applied for triggering MS2 scans. Monoisotopic peak selection was enabled and set to isolate the most abundant peak in peptide mode. Isolated masses were excluded for 60 s after triggering with a mass tolerance window of ±10 ppm, while also excluding isotopes and different charge states of the isolated species. A charge state filter was set for 2-5+ charge states.

The following settings were used for Orbitrap™ MS2 scans. AGC target was set to 1E5 charges, and the maximum IIT was 96 ms. The quadrupole was utilized for isolation with an isolation width of 0.4 Da, and ions were fragmented with HCD at a normalized collision energy of 30%. The resolution was 45k with a defined first mass at 110 m/z.

### Data Analysis

Raw data files were analyzed using Proteome Discoverer 3.1.1.93 (Thermo Scientific) with the SEQUEST HT search engine. The database search was performed against the reference proteomes for *X. laevis* (XenBase v10.1: https://www.xenbase.org), *S. cerevisiae* (S288C: UP000002311), and *H. sapiens* (UP000005640). The search included concatenated target-decoy entries to enable FDR estimation. For methods employing Orbitrap™ MS1 scans, a precursor ion mass tolerance of 20 ppm was used, whereas iTMTproC analyses utilizing ion trap MS1 scans applied a tolerance of 0.7 Da. Similarly, Orbitrap™ MS2 scans used a fragment ion mass tolerance of 0.02 Da, while ion trap MS2 scans were searched with a tolerance of 1 Da. Spectra were searched using full tryptic digestion with LysC and trypsin specificity, allowing up to two missed cleavages. N-ethylmaleimide (NEM) on cysteine (+125.048 Da) and TMTpro on the peptide N-terminus and lysine was set as a static modification (+304.207 Da). Oxidation of methionine (+15.995 Da) was considered a variable modification. MS2 spectra were selected using the Spectrum Selector node, followed by SEQUEST database searching. Percolator was used for post-search validation, with a target-decoy strategy to control the PSM FDR at 1%. Protein-level confidence was assigned by estimating q-values derived from comparisons between target and decoy protein scores across a range of thresholds. Grouping was then performed to consolidate proteins with overlapping or nested peptide identifications, resulting in a parsimonious list of master proteins. Final protein tables were filtered to include only groups with at least one unique peptide and passed 1% protein-level FDR. For analysis of the human-yeast interference standard, only unique yeast peptides were used for quantification to avoid confounding effects from shared razor peptides between human and yeast.

Complement ion quantification was performed using previously described methods without modification^27, 28^. Briefly, the m/z of each complement reporter ion was calculated based on theoretical TMTpro fragmentation, and the observed intensities were extracted. Isotopic impurity correction was applied using a custom R script that integrates tag-specific isotopic impurities and peptide isotopic distributions to resolve an overdetermined system via QR decomposition, yielding corrected complementary ion ratios. This pipeline is described in greater detail elsewhere^27, 28^, and the quantification module is scheduled for inclusion in an upcoming release of Proteome Discoverer. For iTMTproC, PSMs were filtered based on the following criteria: Charge=2, Theoretical m/z<1075, and M_0_/M_1_ isolation error Δm/z ± 0.2. Complement ion intensities from redundant PSMs passing all filters were summed to retain usable signal that would otherwise be discarded. An intensity threshold of 563 ions was used, corresponding to an expected CV of 10% as described above^31^.

For reporter ion quantification, reporter ion intensities were extracted in the Reporter Ion Quantifier node of Proteome Discoverer. For non-RTS MultiNotch MS3 workflows, only PSMs with co-isolation interference below 25%, SPS matches above 45%, and signal-to-noise of at least 129 for 2+ peptides and 265 for 3+ peptides were retained for quantification. This signal threshold matches the ion statistical criteria of iTMTproC with an expected CV of 10%^27, 31^. Protein-level quantification was performed using median reporter ion intensity across all PSMs assigned to each protein group. TMT-MS2 used the same quantification workflow but excluded the SPS match requirement when filtering PSMs.

### Data availability

The MS proteomics data generated in this study has been deposited to the ProteomeXChange Consortium via the PRIDE^55^ partner repository with the dataset identifiers PXD062540.

## Acknowledgement

This work was supported by the National Institutes of Health grant R35GM128813 (MW). We gratefully acknowledge support by the Princeton Catalysis Initiative and Eric and Wendy Schmidt Transformative Technology Fund. We are grateful to Dean Edelman and his colleagues in the Office of the Dean of Research for their support in enabling this study. We gratefully acknowledge support by the American Heart Association predoctoral fellowship 20PRE35220061 (TN) and NSF Graduate Research Fellowship (ERC). We thank Eric Wieschaus and Trudi Schüpbach for useful discussions and suggestions in writing this manuscript. The content of this article is solely the responsibility of the authors and does not necessarily represent the official views of the National Institutes of Health.

## Conflict of Interest

GCM, PMR, and CJ are employees of Thermo Fisher Scientific. Thermo Fisher provides limited support to MW laboratory under a collaborative research agreement with Princeton University. All other authors declare they have no conflicts of interest with the contents of this article.

