## Supporting Information for "Expanding Targeted Instrumentation for Discovery Applications: Complement Reporter Ion Quantification with a Quadrupole-Ion Trap Instrument"

**Table of contents**

- **Figure S1. Scan speed does not affect the conversion factor between charges and pseudo-counts in an ion trap.**
- **Figure S2. Evaluation of ion trap scan speeds for iTMTproC quantification using the human–yeast interference standard.**
- **Figure S3. Comparison of iTMTproC with MultiNotch MS3 and TMTpro-MS2 on the Ascend™.**

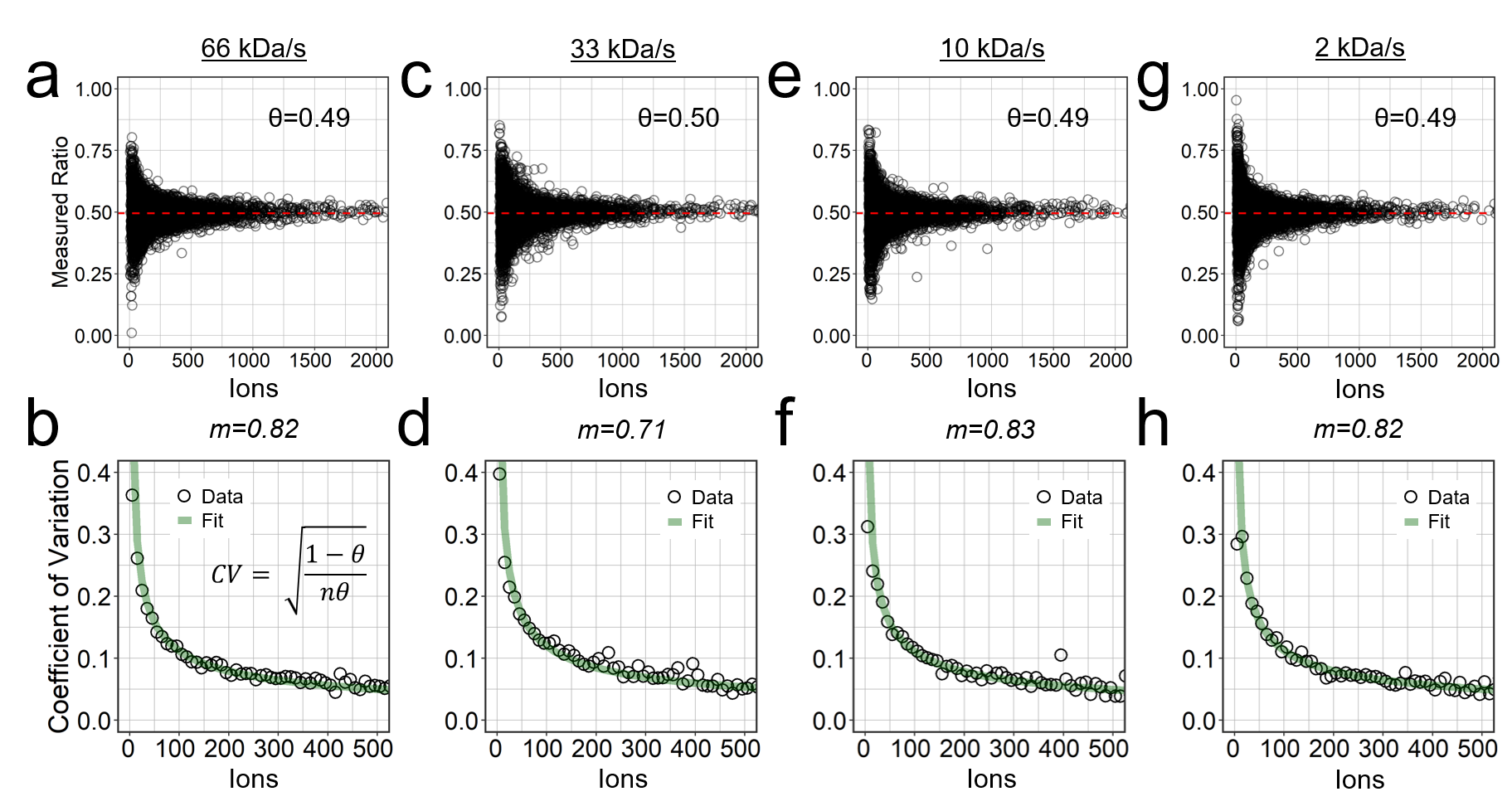

**Figure S1. Scan speed does not affect the conversion factor between charges and pseudo-counts in an ion trap.**

Panels (a, c, e, g) show measured peptide ratios plotted against summed charges at different scan speeds (66, 33, 10, and 2 kDa/s, respectively). The expected ratio (θ) remains consistent across all conditions. Panels (b, d, f, h) display the coefficients of variation (CVs) as a function of charges for each corresponding scan speed. The conversion factor, derived from binomial modeling, remains stable across scan speeds, confirming that—unlike in the Orbitrap™—this factor does not depend on resolution in an ion trap. All data were acquired in a single run, testing all four scan speeds in parallel.

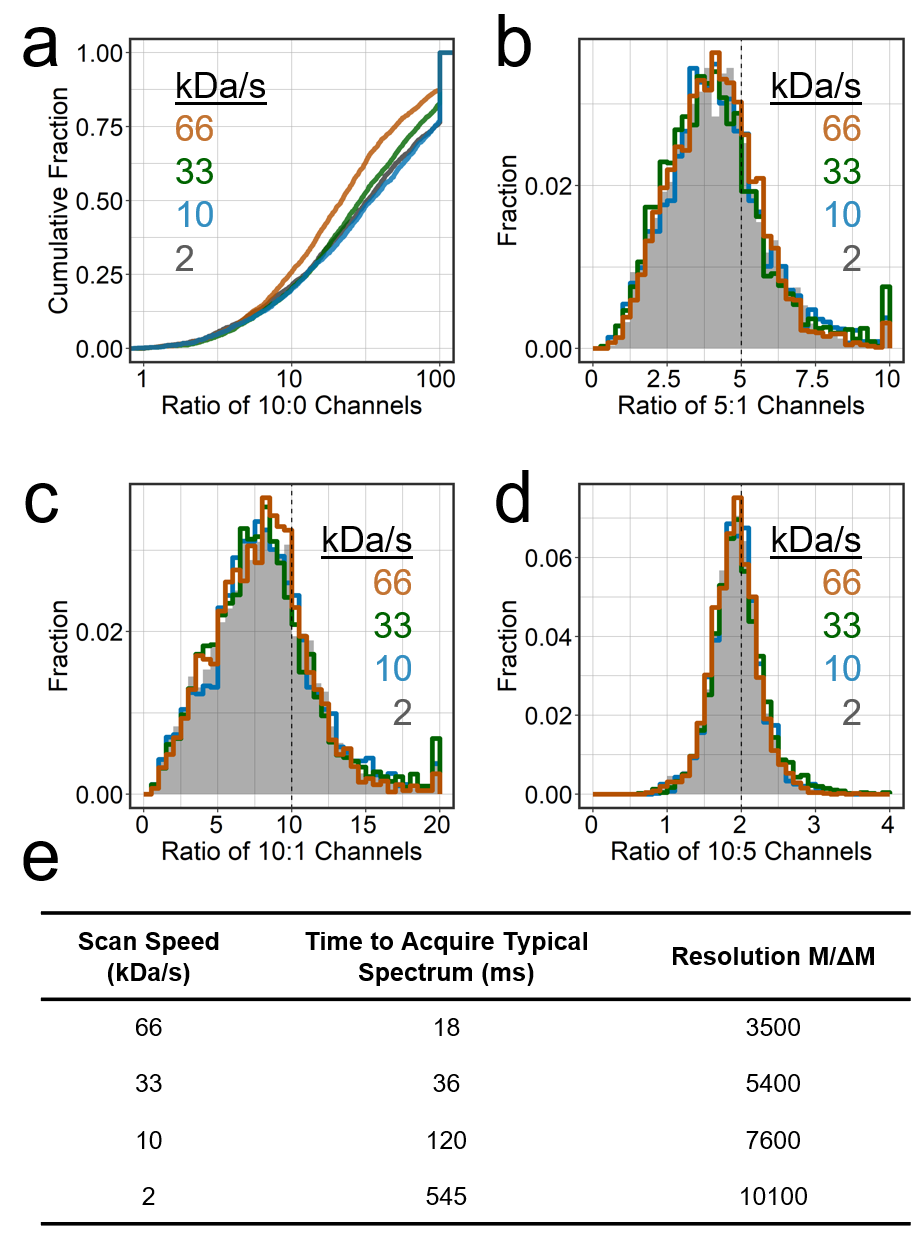

**Figure S2. Evaluation of ion trap scan speeds for iTMTproC quantification using the human–yeast interference standard.**

Four ion trap scan speeds (66, 33, 10, and 2 kDa/s) were tested in a single run to determine the fastest speed that maximizes peptide identifications while maintaining data quality.

(a) Cumulative distribution of measured 10:0 channel ratios. Lower cumulative fractions indicate reduced interference.

(b–d) Histograms of measured ratios for the interference standard: 5:1 (b), 10:1 (c), and 10:5 (d). Results show that scan speed has minimal impact on resolving complement peaks across all speeds tested. However, 66 kDa/s was found to be sensitive to calibration drift and was deemed impractical for routine use.

(e) Summary table of manufacturer-reported scan speed specifications. For each scan speed, the corresponding time to acquire a typical 1200 m/z spectrum and the reported resolution (M/ΔM) are listed. Based on these findings, 33 kDa/s was selected as the standard scan speed for iTMTproC due to its optimal balance of speed, resolution, and robustness.

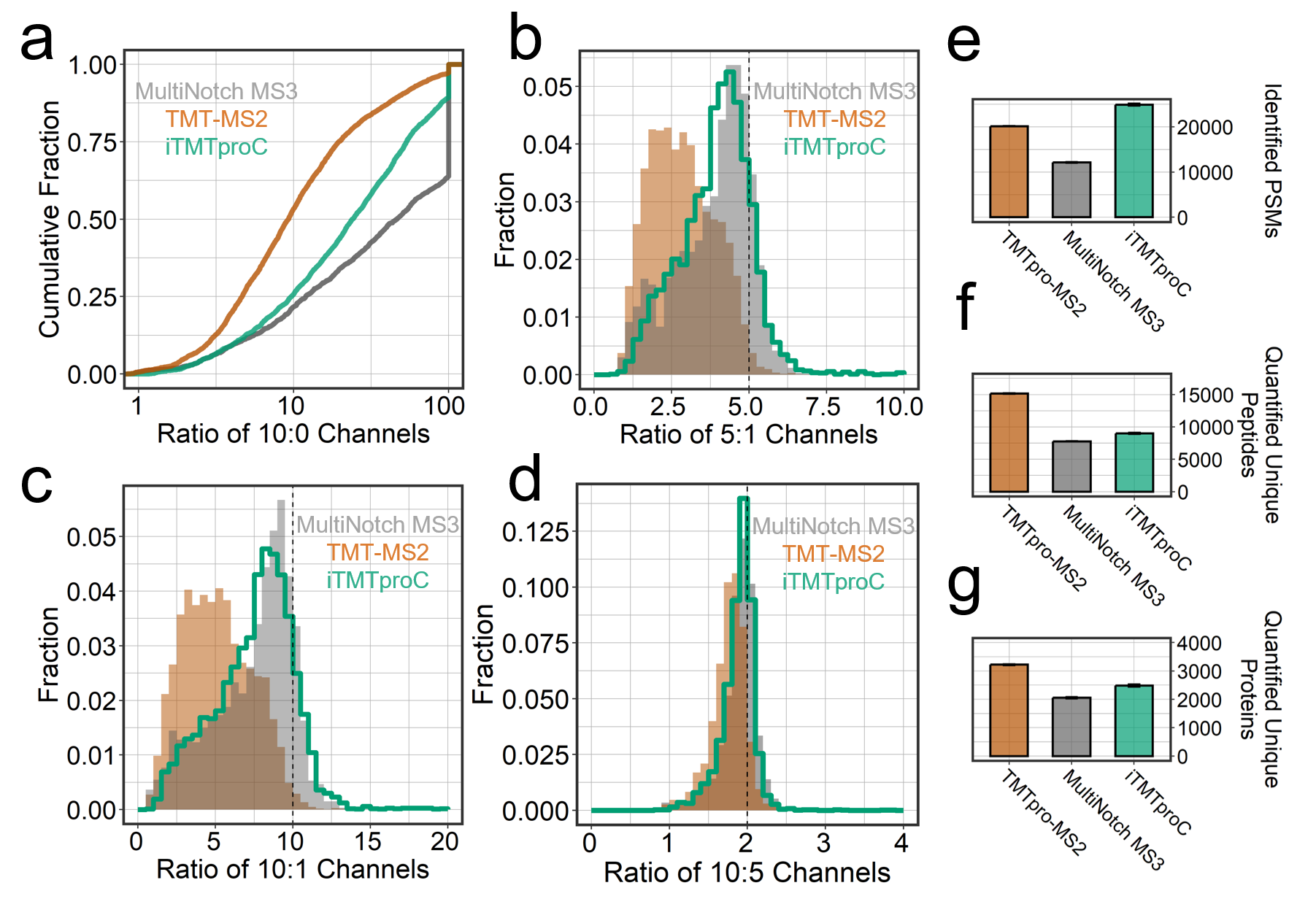

**Figure S3. Comparison of iTMTproC with MultiNotch MS3 and TMTpro-MS2 on the Ascend™.**

(a) Cumulative distribution of the measured 10:0 channel ratios. Since the true 10:0 ratio is infinite, a lower cumulative fraction less than 100 suggests reduced interference. TMTpro-MS2 shows the most distortion, and ion trap TMTproC (iTMTproC) outperforms TMTpro-MS2 but is still behind MultiNotch MS3. Data for iTMTproC, TMTpro-MS2, and MultiNotch MS3 was acquired on an Ascend™, ensuring a fair comparison of all methods using a consistent gradient and an instrument capable of accurately representing all three approaches. The Ascend™ has a brighter beam and more parallelizable injection time than a Fusion™ series MS. However, for FTMS2 methods, it is reasonable to compare this to data quality achieved on an Orbitrap Lumos™.

(b-d) Histograms of measured ratios for 5:1 (b), 10:1 (c), and 10:5 (d) channels. Ratios closer to the expected values indicate less interference and improved accuracy. Further illustrating the trend observed in (b), iTMTproC exhibits comparable interference reduction as MultiNotch MS3 on the same instrument.

(e–g) Peptide and protein identification. Like analyses on independent instruments, iTMTproC yields the highest number of PSMs and quantifies more unique proteins than MultiNotch MS3. As an intermediate approach, iTMTproC offers higher sensitivity than MultiNotch MS3 but slightly lower data quality, while greatly improving data quality over TMTpro-MS2 with a reduction in sensitivity. Error bars are shown and represent the standard error of the mean (n = 3).
